# Local intracerebral Inhibition of IRE1 by MKC8866 sensitizes glioblastoma to irradiation/chemotherapy *in vivo*

**DOI:** 10.1101/841296

**Authors:** Pierre Jean Le Reste, Raphael Pineau, Konstantinos Voutetakis, Juhi Samal, Gwénaële Jégou, Stéphanie Lhomond, Adrienne M. Gorman, Afshin Samali, John B Patterson, Qingping Zeng, Abhay Pandit, Marc Aubry, Nicolas Soriano, Amandine Etcheverry, Aristotelis Chatziioannou, Jean Mosser, Tony Avril, Eric Chevet

## Abstract

Glioblastoma multiforme (GBM) is the most severe primary brain cancer. Despite an aggressive treatment comprising surgical resection and radio/chemotherapy patient’s survival post diagnosis remains short. A limitation for success in finding novel improved therapeutic options for such dismal disease partly lies in the lack of a relevant animal model that accurately recapitulates patient disease and standard of care. In the present study, we have developed a novel immunocompetent GBM model that includes tumor surgery and a radio/chemotherapy regimen resembling the Stupp protocol and we have used this model to test the impact of the pharmacological inhibition of the endoplasmic reticulum (ER) stress sensor IRE1, on treatment efficacy.

## Introduction

Glioblastoma multiforme (GBM) is the most severe primary brain cancer and represents more than 15% of primary brain tumors. Despite an aggressive treatment comprising surgical resection and a combination of irradiation and chemotherapy (“Stupp protocol”), patient’s survival post-diagnosis remains short with a median overall survival of 15 months [1]. The lack of efficacy of the current treatments is mostly due to the intratumor heterogeneity at diagnosis with the presence of multiple driver alterations [2], as well as to different tumor cell types that exhibit various sensitivity to anti-cancer agents. Finally, it is also due to the diffuse nature of GBM tumors that precludes from performing complete resection. Another limitation for the discovery of new improved therapeutic approaches is the lack of a relevant animal model that extensively recapitulates current GBM patient disease and standard of care.

In the past couple of years, we have demonstrated that the Unfolded Protein Response (UPR) plays an instrumental role in GBM development [3-8]. More precisely, we have shown that IRE1, the most conserved UPR sensor, signals in tumor cells to remodel the tumor microenvironment, to favor tumor growth and most likely to alter the response to treatment. We demonstrated that the IRE1/XBP1s (spliced XBP1) axis exhibits pro-oncogenic properties, whereas Regulated IRE1 Dependent Decay of RNA (RIDD) dampens tumor angiogenesis and tumor cell migration/invasion. This observation was then used to stratify GBM patients, where we showed that survival of patients with tumors exhibiting high XBP1s and low RIDD was significantly reduced compared to that of patients with low XBP1s and high RIDD tumors [8]. These observations point toward the IRE1/XBP1s axis as a potentially relevant therapeutic target which, when inhibited in XBP1s high tumors would i) slow down tumor growth by impairing the pro-tumoral remodeling of the tumor stroma and ii) sensitize tumor cells to the current treatments [8].

To further test the potential impact of the pharmacological targeting of IRE1/XBP1, we propose to use an IRE1 RNase inhibitor, MKC8866 [9], already shown to be effective in other cancer models [10], in preclinical models of GBM. However, the current preclinical GBM models poorly recapitulate the clinical handling of GBM patients, therefore would not be appropriate to test the therapeutic potential of IRE1 inhibition. In the present study, we developed a novel immunocompetent GBM model that involved tumor surgery and a radio/chemotherapy regimen, resembling the Stupp protocol. Furthermore, we used this unique model to test the therapeutic relevance of IRE1 inhibition on tumor microenvironment and on treatment efficacy.

## Materials and methods

### Reagents

All reagents were purchased from Sigma-Aldrich (*St Quentin Fallavier, France*), unless otherwise stated. For immunohistochemistry staining, primary antibodies against vimentin (rabbit monoclonal, used at 1 in 250 dilution) was obtained from Abcam (*Paris, France*); CD31 (rabbit polyclonal, used at 1 in 50 dilution) from Bioss Inc. (*Cliniscience, Nanterre, France*); and against IBA-1 (rabbit polyclonal, used at 1 in 50 dilution) from Wako (*Sobioda, Montbonnot, France*). For Western blot experiments, primary antibodies against ATF4 and phospho-eIF2alpha were purchased from Cell Signaling Technology (*Ozyme, St Quentin-en-Yvelines, France*), anti-Calnexin (CNX) antibodies were a kind gift from JJM Bergeron (McGill University, Montreal, Qc, Canada).

### Cell lines

The GL261 murine glioblastoma cell line was a kind gift from Dr Clavreul (University of Angers, France). These cells were transfected with the plasmid pGL4-Luc and the luciferase-expressing cells stably selected using G418 (the selected line was polyclonal and renamed GL261-Luc). The cells were cultured in Dulbecco’s Modified Eagle Medium *(DMEM, Life Technologies, Thermo Fisher Scientific, Courtaboeuf, France)* with the addition of 10% FBS *(Thermo Fisher Scientific)*, and 100 μg/ml of G418 *(Thermo Fisher Scientific)*. The cells were grown in a humidified incubator at 37°C with 5% CO_2_. The human U251 GBM cells were grown in DMEM containing 10% FBS.

### Genome analyses

Whole Genome Sequencing of the GL261 cell line was performed by Integragen (https://www.integragen.com/fr/). PCR free Libraries were prepared with NEBNext Ultra II DNA Library Prep Kit according to the supplier recommendations. Briefly the key stages of this protocol comprise a specific double strand gDNA quantification, a fragmentation (300 ng of input High molecular weight gDNA) using sonication method to obtain approximately 400 bp medium pieces, and finally, paired-end adaptor oligonucleotides (xGen TS-LT Adapter Duplexes from IDT) are ligated on repaired A tailed fragments then purified for direct sequencing without PCR step. DNA PCR free library was sequenced on Paired End 150 bp run on the Illumina HiSeq4000. Image analysis and base calling were performed using Illumina Real Time Analysis (RTA) Pipeline version 2.7.7 with default parameters. Sequence reads were mapped to the mouse genome build (mm10) by using the Burrows-Wheeler Aligner (BWA) tool. The coverage was 25X for 75% of the mouse genome with a mean depth of 33X. The duplicated reads (e.g. paired-end reads in which the insert DNA molecule showed identical start and end locations in the mouse genome) were removed (samba tools). Variant calling, allowing the identification of genetic alterations, as well as SNV (Single Nucleotide Variation) small insertions/deletions (up to 20 bp), was performed via the Broad Institute’s GATK Haplotype Caller GVCF tool (3.7) for constitutional DNA. Finally, an in-house post-processing to filter out candidate germline mutations that are more consistent with artifacts was applied (Mapping Quality, Base Quality, Strand Bias, Variant Quality). Ensembl VEP (Variant Effect Predictor, release 87) program processed variants for further annotation. This tool annotates variants, determines the effect on relevant transcripts and proteins, and predicts the functional consequences of variants. It takes into account data available in dbSNP (dbSNP146). Other information like quality score, homozygote/heterozygote status, count of variant allele reads were also reported. We detected copy-number variants and allelic imbalances in GL261 using the Control-FREEC algorithm (FREEC-11.5) with the C57BL_6.bam file downloaded from the SANGER FTP server (ftp://ftp-mouse.sanger.ac.uk/current_bams/) as a non-tumor reference sample.

### Transcriptome analyses

In order to quantify the transcript-level abundances of implanted GL261-derived tumors for the seven experimental conditions (NR:no resected tumors; R:resected tumors without treatment; MKC:treatment with MKC8866 inhibitor; STUPP protocol; STUPP_MKC: STUPP protocol including MKC8866 inhibitor; PARENTAL:GL261 parental; STUPP_GELL: STUPP protocol including gel implantation lacking MKC8866 inhibitor) RNA sequencing was performed by Integragen (https://www.integragen.com/fr/). RNAseq libraries were prepared with the NEBNext Ultra II Directional RNA Library Prep Kit for the Illumina protocol according to the supplier recommendations. Briefly the key stages of this protocol included the purification of PolyA containing mRNA molecules using poly-T oligo attached magnetic beads from 1μg total RNA (via the Magnetic mRNA Isolation Kit from NEB), a fragmentation step using divalent cations under elevated temperature to obtain approximately 300bp pieces, double strand cDNA synthesis and finally Illumina adapters ligation and cDNA library amplification by PCR for sequencing. Sequencing was then carried out on paired-end 75 bases of Illumina HiSeq4000. Image analysis and base calling were performed using Illumina Real Time Analysis (RTA) Pipeline version 2.7.7 with default parameters. Starting from the FASTQ files, qualitative evaluation of reads was executed using the FastQC software (http://www.bioinformatics.babraham.ac.uk/projects/fastqc/) with 95% of bases scoring Q30 and above. Transcript-level quantification data were produced using Salmon (v.1.0.0) program [11], installed in Linux server environment (Ubuntu 16.04.6 LTS) by performing mapping of sequence reads to the reference mouse transcriptome (Release M23 - GRCm38.p6) downloaded from GENCODE (https://www.gencodegenes.org/mouse/release_M23.html). For the building of transcriptome index the following command was executed: “salmon index --gencode -t gencode.vM23.transcripts.mouse.fa.gz -i gencode.vM23_salmon_1.0.0” and the provided index was used for the quantification step. For the salmon quantification process, a simple shell script was created, defining the directory address of transcriptome index and the library type (-l ISR; “fr-firststrand”; R1-reverse strand and R2-transcript strand) of RNAseq data while the --validateMappings and GC bias correction (--gcBias) setting was chosen. Estimated counts per transcript as well as effective transcript lengths which summarize bias effects were calculated. After running Salmon, the tximport R/Bioconductor package [12] was used to summarize the estimated transcript abundances to gene-level count matrices correcting for changes in transcript length across samples. For this reason, the tximport argument “countsFromAbundance” was assigned to “lengthScaledTPM” value and bias corrected gene-level count matrix (txi$counts) was produced. Using the annotation R package “org.Mm.eg.db” [13] the Ensembl Gene IDs mapped to external Gene Symbols and with the Bioconductor differential gene expression method, edgeR, [14] a DGEList object was created. After count matrix filtering by keeping only genes expressed in at least one condition, TMM normalization was executed and log_2_CPM expression values were calculated using the cpm() function of edgeR package. The log_2_CPM gene expression matrix was used for gene-level exploratory data analysis (EDA) with visual quality assessment providing insight into the possible relationships between the samples. For the conversion of mouse gene symbols (mgi_symbol) to human gene symbols (hgnc_symbol) the getLDS() function of biomaRt R package was used retrieving homologous genes after linking of “mmusculus_gene_ensembl” and “hsapiens_gene_ensembl” datasets. Mice do not have a structural homolog for CXCL8 and MMP1 human genes and Cxcl1 and Mmp1a & Mmp1b were used instead, because of their functional similarity [15-17]. The expression profile of U87-MG human glioblastoma cells (WT-wild type) bearing also, different IRE1 mutants (A414T, P336L, Q780Stop, S769F) and a dominant negative form of IRE1 (DN), under normal and stress conditions (tn; tunicamycin treated), deposited in GEO (Gene Expression Omnibus) [18] (GSE107859 accession number) was compared with the gene expression profile of GL261 parental cell line after cross-platform harmonization, and evaluation of IRE1α signaling activity with use of IRE1α signature (XBP1s and RIDD component mapping) as described in Lhomond et al. 2018 [19]. For the cross-platform normalization process, the Shambhala algorithm [20] was performed in R package HARMONY (https://github.com/oncobox-admin/harmony) for a set of 6546 genes in common between the two platforms (Affymetrix microarrays and RNA sequencing). For the exploration of relationships between the samples cluster dendrograms were built using R package ComplexHeatmap [21] and the statistical significance of hierarchical clustering was assessed with bootstrapping method using the R package pvclust [22]. Sample distance matrices to assess the overall similarity between samples were produced using the R function pheatmap() of R pheatmap package [23] with calculation of Euclidean distance between samples and minimization of the total within-cluster variance by Ward’s minimum variance method (ward.D2) [24]. Finally, principal component analysis (PCA) on harmonized gene expression data was conducted with prcomp() function including centering and scaling to unit variance while the visual projection of principal components was based on ggbiplot() function, an implementation of the biplot using the R package ggplot2 [25].

### Cell culture and treatments

For survival assays, GL261 cells were cultured in 96-well plates at 5,000 cells/well in the presence of increasing amounts of temozolomide (TMZ) (0 to 1000 µM), MKC8866 (0 to 200 µM) with two treatments of radiation at 5 Gy (at day 1 and day 3). After 5 days (for TMZ) or 6 days (for MKC8866) of culture, 20 µl of the WST1 reagent were added to each well. After 4 hours at 37°C, optical densities (OD) were analyzed using spectrophotometry at 450 nm and 595 nm. Specific OD were given by the difference between the OD observed at 450 nm and the OD at 595 nm. Cell viability was calculated by the ratio of the specific OD observes with cells incubated in the presence of different concentration of TMZ and the specific OD observed with cells cultured in medium alone, this information allowed us to determine IC50 values using Prism software (GraphPad). Each experimental point represents at least a triplicate. For protein and RNA extractions, GL261 cells were seeded at a density of 10^6^ cells per dish (6 cm diameter), with two dishes per time point. Cells were treated as described above for 2, 4, 8 and 24 hours. Each experimental point represents at least a triplicate. For protein extraction, cells were lysed using 100 µl of lysis buffer (30 mM Tris-HCl, pH 7.5, 150 mM NaCl, 1.5% CHAPS and Complete™) per plate. Clarified lysates were complemented with 20 µl of 5 times reducing Laemmli sample buffer prior heat-denaturation and used in Western blot experiments with the indicated antibodies (at 1/1000 dilutions). For RNA preparation, cells were lyzed using 1ml of Trizol reagent per plate and RNA was extracted according to the manufacturer’s instructions (*Thermo Fisher Scientific*) and PCR analyses performed using previously described protocols [11].

### Tumor cell orthotopic implantation

Tumor cells (GL261-Luc) were implanted into the brain of immunocompetent C57BL/6rJ, 8 weeks old male mice *(Janvier, Laval, France)*. All animal procedures met the European Community Directive guidelines (Agreement B35-238-40 Biosit Rennes, France/ No DIR 13480) and were approved by the local ethics committee and ensuring the breeding and the daily monitoring of the animals in the best conditions of well-being according to the law and the rule of 3R (Reduce-Refine-Replace). GL261-Luc cells were implanted in the mouse brain by intracerebral injection followed by tumor growth monitoring using bioluminescence. The mice were anesthetized intraperitoneally (i.p.) and then fixed on a stereotactic frame. This framework makes it possible to manipulate the brains of living animals, and to reach isolated areas of the brain precisely relative to markings visible to the naked eye through the use of three-dimensional coordinates. After incising the scalp, the stereotaxic coordinates were calculated for injection of tumor cells into a specific point of the brain, and reproducible for all the mice used. In the study, the tumor cells (2.5 × 10^4^ cells per mice in 1 μL) were injected at 2.2 mm to the left of the bregma and 3.2 mm deep to perform the implantation at the level of the striatum.

### Bioluminescence

Mice were assessed with *in vivo* bioluminescence imaging every 3 days, for the first week, and weekly onward for tumor progression and followed for signs of neurologic deterioration daily. Mice were injected i.p. with 100 μl of luciferin (*Promega, Charbonnières-les-Bains, France*). The luciferin was allowed to circulate for 10 min before the mice were anesthetized with a mix of O_2_ and isoflurane (2.5%). Mice that showed an increase in tumor burden based on imaging during the first week after tumor implantation were included in the study. Bioluminescence analysis was used to determine the window of resection (day 14).

### Resection

The mice were anesthetized by the intraperitoneal route with 90 μl of anesthetic (1.5 mg/kg of ketamine and 150 μg/kg of xylazine). Then they were attached to the stereotaxic frame and the scalp was incised to expose the convexity of the skull and the injection point. The opening of the bone was performed with thin forceps allowing minimal craniectomy. Visualization of the tumor was enhanced by using an injection of intravenous fluorescein at the beginning of the procedure. Thanks to a UV source, this fluorophore makes possible to visualize the zones of rupture of blood-brain barrier characteristic of glioblastoma. This visual aid makes it possible to delineate the tumor borders and thus minimizes the lesions on healthy parenchyma. The resection was performed using a surgical microscope, and the tumor was aspirated with the help of a small suction. The extent of resection was mainly driven by the depth of the tumor. Once the resection was performed, the hole left by the tumor was filled with a fibrin-collagen gel containing the inhibitor of interest. The craniectomy was closed with the help of a teflon patch (Neuropatch^®^) slid under the skull, fixed by the application of biocompatible acrylic glue. This step allows maintenance of intracranial pressure and prevents tumors from developing in the subcutaneous space. Finally, the skin was sewn with non-resorbable monofilament sutures at the end of the procedure (see **Movie S1**).

### Stupp-like treatment applied to mice

Three days after the resection, we proceeded to apply the Stupp protocol, consisting in two steps including (i) concomitant radio/chemotherapy: mice were anesthetized with Isoflurane, and placed in a lead dome to cover the entire animal to prevent ionizing radiation from reaching the healthy tissues of the animals. A hole of a few centimeters was made in the lead dome, corresponding to the resection area of the tumor, in order to irradiate only the area of interest. The animals were subjected to a radiation of 5 times 2 Gy 3 days after resection along with 25 mg/kg/day of TMZ delivered intraperitoneally followed by (ii) stand-alone chemotherapy. In the latter procedure animals were subjected to the standard procedure of chemotherapy, namely 30-50 mg/kg of TMZ on 5 days with 2 or 3 days of rest between each treatment. Survival was measured as the time between implantation and sacrifice, which was performed by cervical dislocation in case of critical clinical signs or loss of weight >15%.

### Immunohistochemistry

The immunochemistry experiments were performed on 5 µm tumor sections which were incubated at room temperature for 1 hour with primary antibodies. Immunostaining was performed using BenchMarkXT-Ventana Medical Systems with kit OMNIMAP (system “biotin-free” using multimer technology) with antigen retrieval for all (Tris/borate/EDTA pH 8). To perform the analysis, glass slides were converted on to digital slides with the scanner Nanozoomer 2.0-RS Hamamatsu.

### Manufacturing of the gel implant containing the IRE1 RNase inhibitor MKC8866 and cell-based testing

Human fibrinogen was dissolved in deionized water at a concentration of 30 mg/ml followed by dialysis against tris-buffered saline (TBS) overnight. MKC8866 was resuspended in the fibrinogen component of fibrin microgels at a final concentration of 10 and 100 µM. The gel-forming fraction containing MKC8866 was then dispensed as 7 μl droplets on a hydrophobic surface (commercial Teflon^®^ tape) to create a micro-droplet. The gel fractions were then crosslinked by human thrombin (40 U/ml) and incubated for 20 minutes at 37 °C for stable cross-linking. To test the gel in vitro, U251 cells were seeded in 12-well plates at 10,000 cells/well or 20,000 cells/well. After 24h, the cells were cultivated in the absence (no gel) or presence of fibrin microgel containing different concentrations of MKC8866 (0 to 100 µM). After seven days, the cells were treated for 5h with tunicamycin at 5 µg/mL to activate IRE1 pathway and XBP1 RNA splicing. They were then lyzed using 0.5 ml of Trizol reagent per well and RNA was extracted according to the manufacturer’s instructions (Thermo Fisher Scientific). PCR was performed for total XBP1 splicing, and the resulting amplified DNA products were resolved using a 4% agarose gel. The upper band (unspliced XBP1) and lower band (spliced XBP1) were then quantified using the ImageJ software and the ratio between the unspliced (U) and spliced (S) forms of XBP1 were calculated by the following formula: S/(U+S) *100. For in vivo studies, gel implantation was carried out at the moment of the surgery. Briefly preformed microgels were deposited in the resection cavity prior to implanting a teflon plate and sewing the skin.

## Results

### Qualification of GL261 murine GBM as a relevant model for in vivo studies

Since we previously showed that activation of IRE1 signaling (and in particular of the IRE1/XBP1 arm) was linked to GBM aggressiveness by promoting tumor-supporting immune cells, angiogenesis and migration/invasion of the tumor cells [8], we reasoned that a relevant *in vivo* mouse model should i) use GBM cells exhibiting high IRE1 activity, ii) be immunocompetent and iii) be subjected to a treatment similar to the standard of care in human. As such our first objective was to identify a mouse GBM cell line that could be considered as relevant. To this end we investigated the activation status of the IRE1 pathway in the mouse GL261 line. We compared the mouse line to the human U87 line expressing various mutants of IRE1 and treated or not with the ER stressor tunicamycin (**Figure 1A**) [8]. A principal component analysis revealed that GL261 co-clustered with U87, overexpressing the wild type IRE1 or the p336L mutant (in which IRE1 is constitutively activated), treated or not with tunicamycin, thereby indicating that the IRE1 pathway might be constitutively activated in GL261 cells as previously observed in other cell lines [10]. This result was confirmed by the use of the IRE1 activity signature [8], and also showed, using hierarchical clustering, that IRE1 was constitutively active in GL261 (**Figure 1B, C**). To further document the genetic characteristics of the GL261 line, we focused on 86 GBM related genes (including variants, **Tables S1-S3**), including 79 genes recently reported [13], and 7 additional genes (two Stupp protocol response predictors *Mgmt* and *Dgki*; the canonical Endoplasmic Reticulum stress sensors *Eif2ak3* (*Perk*), *Atf6* and *Ern1* (*Ire1*); the genes *Pi3k* and *Tert* frequently mutated in IDH-wild type GBM). We found that 17 genes were harboring one or more mutations with a predicted functional impact (**Table 1**). Some of those mutated genes are known oncogenes (*Kras, Met, Pi3kca*) or tumor suppressor genes (*Nf1* and *Trp53*). Cancer genes were also affected by DNA gains (*Egfr, Mdm2, Pdgfra* for instance) or losses (*Cdkn2c* and *Pten*). RNAseq of the parental GL261 cells further indicated that oncogenic mutations were also present and sometimes enriched at the mRNA level in those cells (*Kras, Nf1, Pik3ca* and *Trp53* for instance, **Table 1, Figure 1D, Tables S1-S3**), interestingly the Kras mutation is found in 2% of the human GBM tumors and the pathway is activated in more than 50% of the tumors [2, 14]. These results showed that GL261 cells express known GBM cancer driver genes, can be used in syngeneic models (in C57BL/6 mice), exhibit a high basal IRE1 signaling and therefore represent a suitable cell line for the development of an *in vivo* model relevant to test the impact of IRE1 inhibition as an adjuvant therapeutic strategy.

**Table 1:**
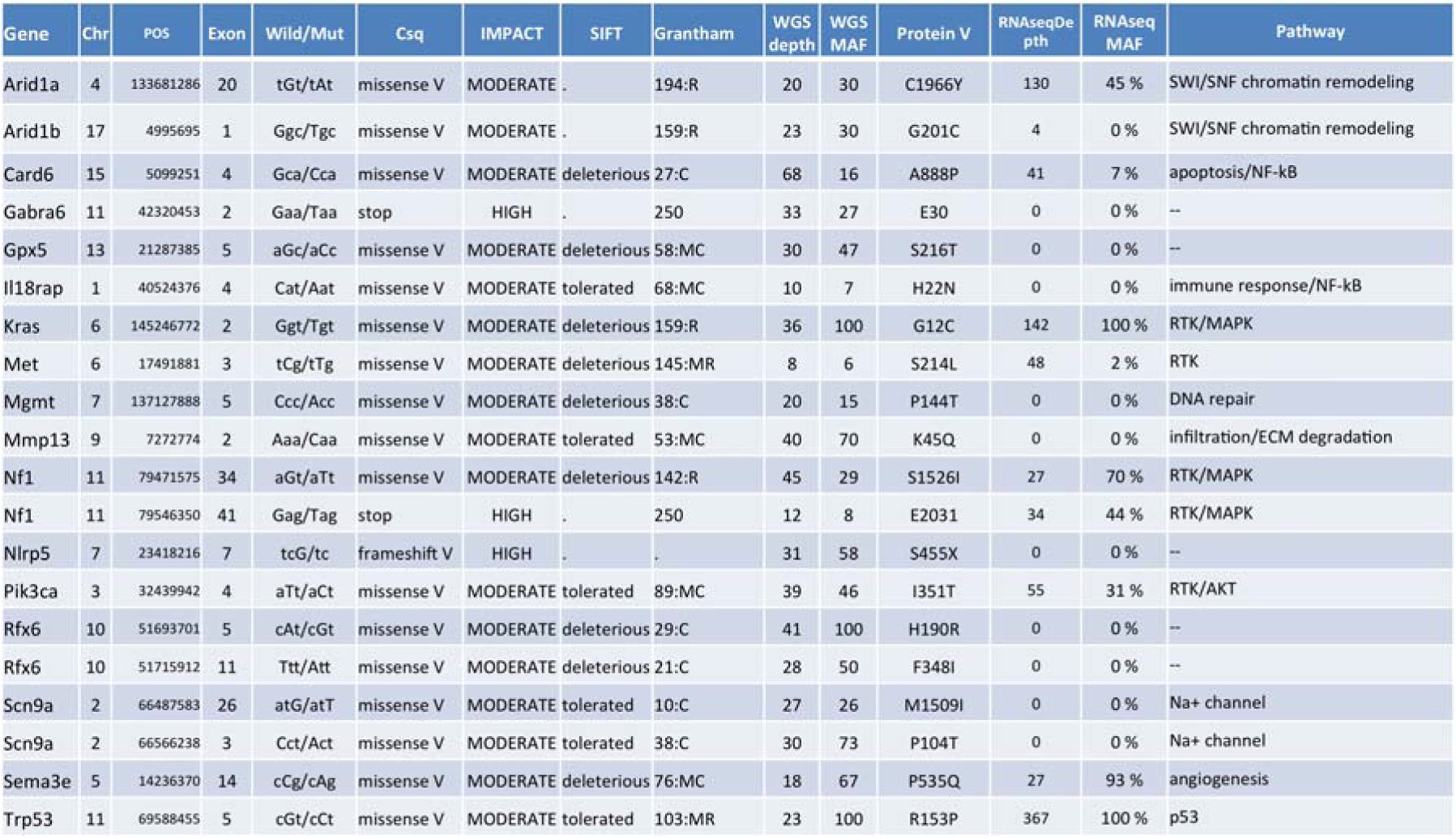
GBM related genes harboring one or more mutations with a predicted functional impact using Ensembl Variant Effect Predictor. Csq: predicted effect that the allele of the variant may have on the transcript. IMPACT: a subjective classification of the severity of the variant consequence, based on agreement with SNPEff (HIGH/MODERATE/LOW).

**Figure 1:**
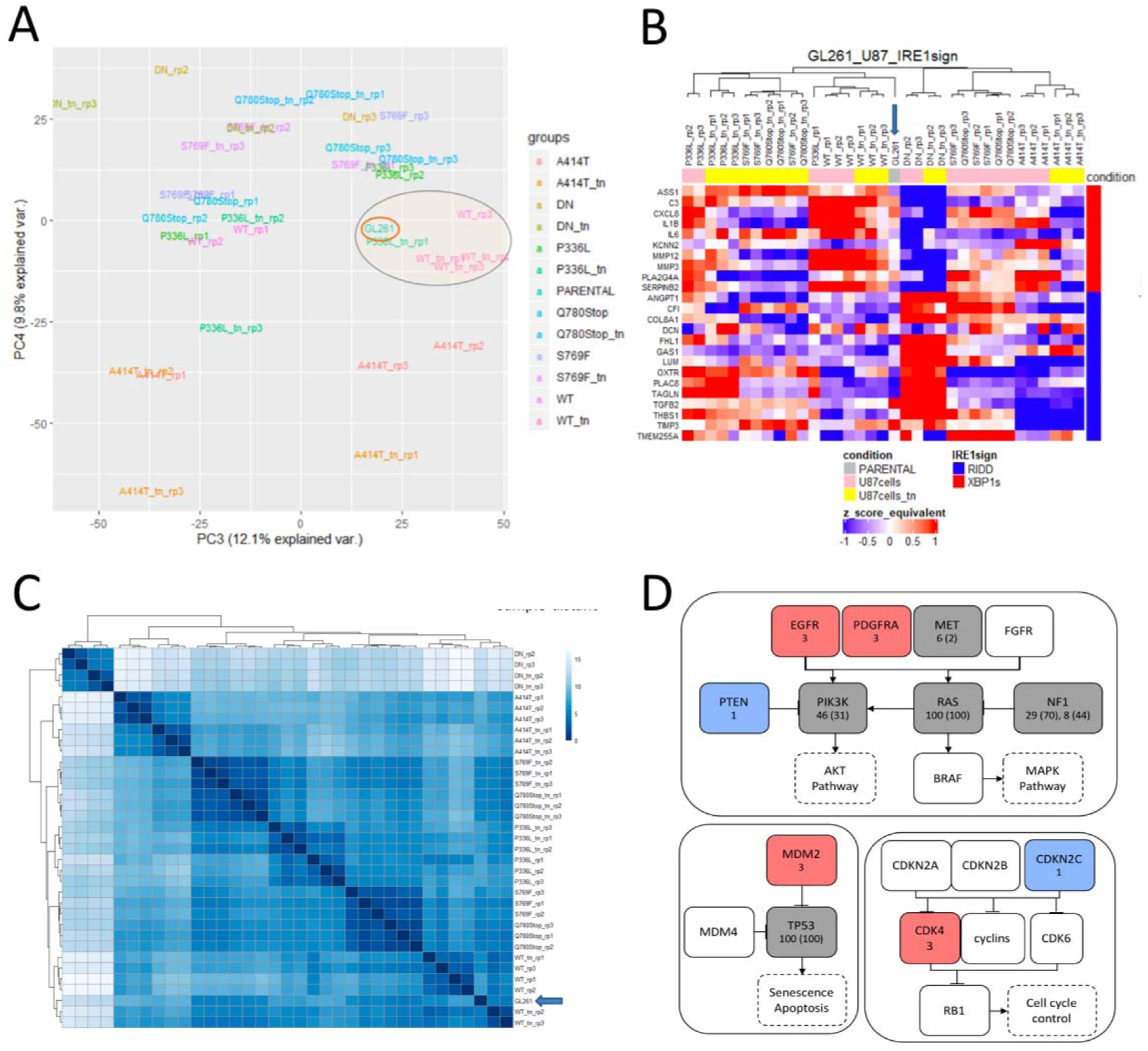
Selection of GL261 as a relevant model for testing the relevance of IRE1 inhibition in GBM. **A)** Biplot projection of PC3 and PC4 components of PCA analysis on harmonized gene expression data of GL261 parental and U87 WT, DN and mutant clones under normal and stress conditions. **B)** Hierarchical clustering heatmap of GL261 parental and U87 WT, DN and mutant cell lines with (tn) and without stress based on the harmonized gene expression profile of IRE1 signature. Pearson correlation coefficient was used for sample distance correlation and the average agglomeration method was used for sample clustering. U87 cells in normal and stress conditions are represented with pink and yellow bars to the top of the heatmap while the GL261 parental cell line is represented with the grey box. **C)** Sample distance matrix of GL261 parental and U87 WT, DN and mutant cell lines under normal and stress conditions, based on the harmonized gene expression profile of IRE1 signature using the euclidean distance between samples. For sample hierarchical clustering Ward’s minimum variance method (ward.D2) was used. **D)** Sequence mutations and copy number changes are depicted as grey boxes for mutations, red boxes for gain of DNA copy and blue boxes for loss of DNA copy. Mutation allele frequency in the gene and the transcript (in parentheses) or DNA copy number are indicated under the corresponding gene name.

### Novel preclinical GBM mouse model including surgery, irradiation and chemotherapy

Due to lack of a murine preclinical model recapitulating GBM patients’ handling, we sought to develop a novel model that would comprise surgical resection combined with radio/chemotherapy, comparable to the clinical setup (**Figure 2A**, human vs. mouse). We designed an experimental approach relying on the mouse GBM line GL261 that we modified for stable expression of luciferase, in order to allow live tumor imaging. These cells were then orthotopically injected in C57BL/6 mouse brain and tumors allowed to grow for 12 to 13 days. At this stage, tumors were surgically resected using fluorescein as a tumor tracer (**Movie S1**) and the mice were allowed to recover from surgery for 1 week before undergoing Stupp-like treatment (**Figure 2B**). The Stupp-like protocol included radiation (2 Gy/day, 5 exposures) combined with chemotherapy (TMZ, 25 mg/kg, 5 applications) and followed by TMZ treatment alone (30-50 mg/g, 5 applications followed by 2 days without treatment for 4 weeks).

**Figure 2:**
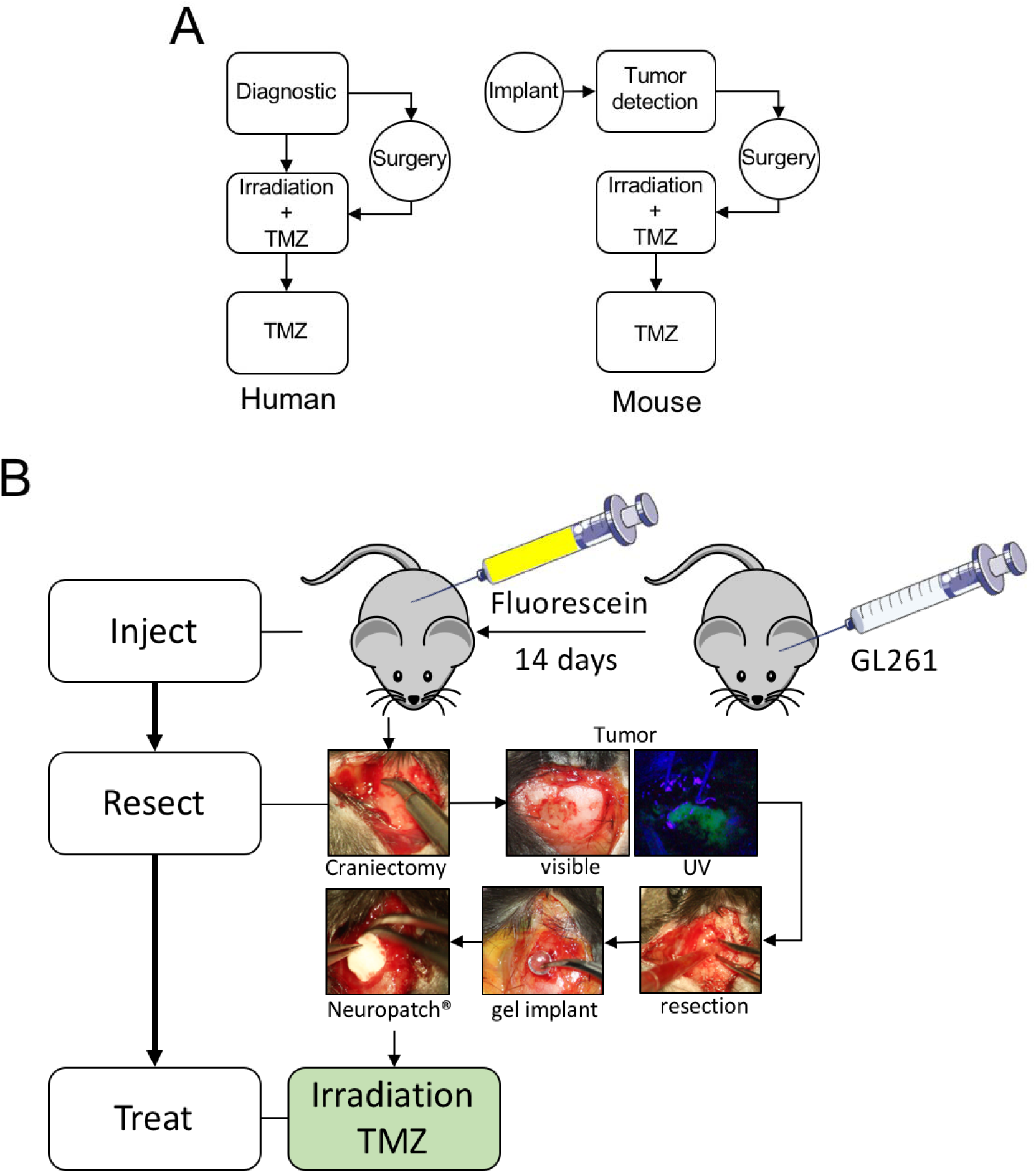
Preclinical development of a novel GBM model in immunocompetent mice. This model comprises surgical resection followed by radio/chemotherapy that recapitulates the current GBM patient standard of care (Stupp protocol) in mice. **A)** Schematic representation of the experimental design and comparison between human and the mouse model. **B)** Schematic representation of the mouse model with the 3 major steps including “Injection” of cells (orthotopic) and fluorescein 14 days later; “Resection” of tumors. The skull was removed and the tumor contour was defined after injecting fluorescein and applying UV source on the tumor site. After aspiration of the tumor material, the resection cavity was filled with hemostatic materials before implanting a teflon prosthesis (replacing the removed bone) and the skin sewn; and “treatment” combining irradiation and chemotherapy (temozolomide/TMZ) – the surgical procedure is also reported in Movie S1.

Following the completion of Stupp-like procedure, described in **Figure 2**, mice were sacrificed at first clinical signs. Brains were collected, fixed in paraformaldehyde (PFA 4%), embedded in paraffin prior to immunohistochemical analysis using anti-vimentin antibodies. This revealed the impact of surgery on tumor size and aggressiveness (**Figure 3A**), as well as the effects of irradiation/TMZ treatment, which appeared to reduce tumor size and ability to infiltrate surrounding parenchyma (**Figure 3A**, SIT). Mouse survival was then evaluated upon these treatments (**Figure 3B**), and showed that while surgery did not have any significant impact, Stupp-like treatment doubled mouse survival, thereby recapitulating observations made in patients following this protocol. Moreover, we analyzed the impact of surgery on tumor angiogenesis by staining the recurring tumor sections with anti-CD31 antibodies (**Figure 3C**). Quantitation of the staining revealed that there was no significant difference in blood vessel density between the non-resected and resected conditions (**Figure 3D**), but more work should be carried out to provide an in depth analysis of the nature of the blood vessels analyzed in both conditions. Furthermore, we analyzed the ability of tumor cells to invade the neighboring parenchyma in primary or post-surgery recurring tumors, by analyzing the vimentin staining (**Figure 3E**). Quantitation of this staining revealed that surgical resection of tumors enhanced the capacity of tumor cells to infiltrate non-tumor tissue (**Figure 3F**).

**Figure 3:**
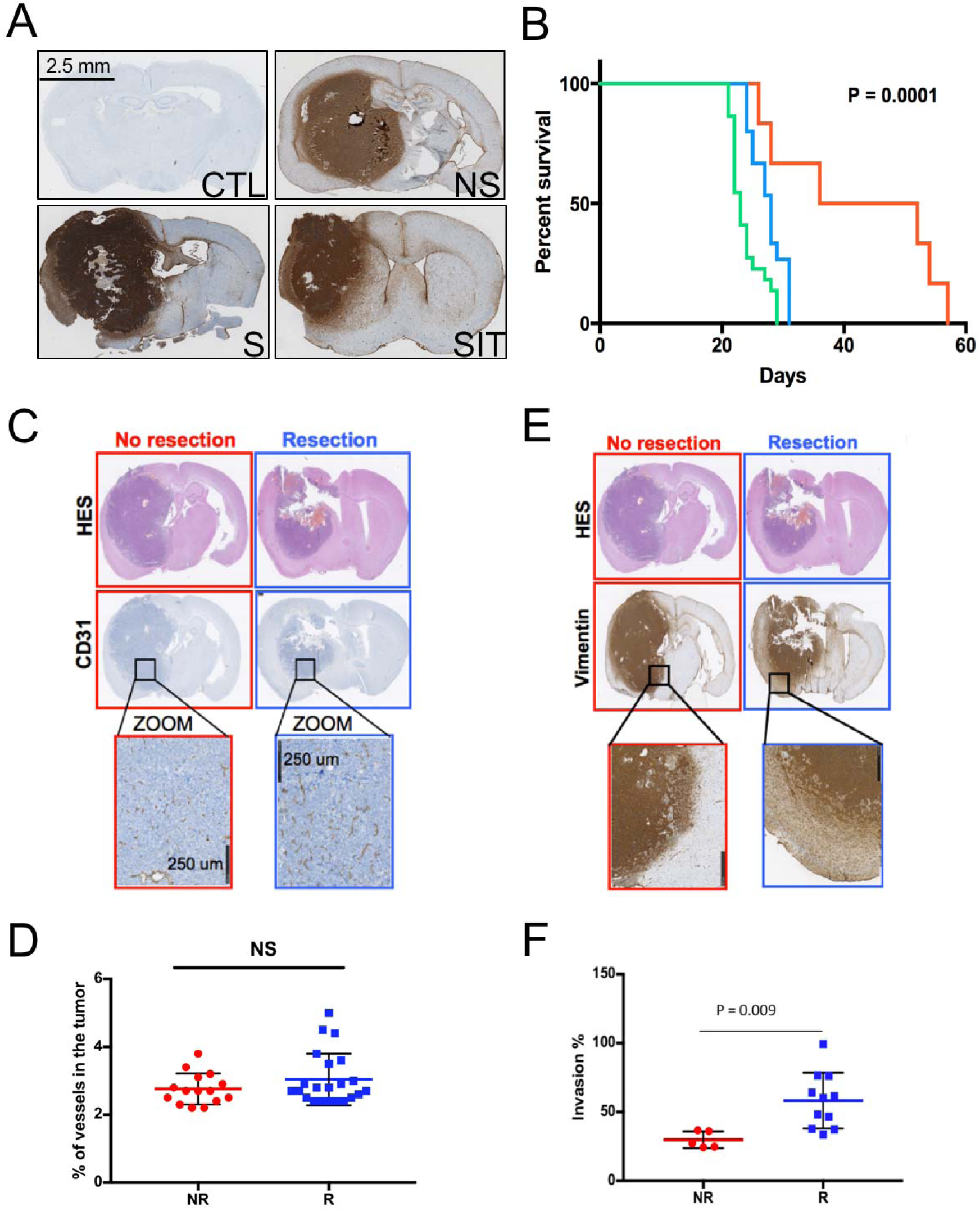
Development and analysis of the establishment of a Stupp-like protocol in mice. **A)** At sacrifice, brains were collected, fixed in PFA, embedded in paraffin prior to be analyzed using immunohistochemistry using anti-vimentin antibodies (CTR: no injection; NS: no surgery; S: surgery only; SIT:surgery/irradiation/TMZ). **B)** Kaplan-Meier representation of mouse survival under this regiment (green: non-resected, blue: surgical resection, orange: surgical resection + radio/chemotherapy). **C-D)** Immunohistochemical analysis and quantitation of tumor angiogenesis post-resection. **E-F)** Immunohistochemical analysis and quantitation of tumor infiltration/invasion post-resection.

### Impact of IRE1 RNase inhibitor MKC8866 in vitro and in vivo

Once this model was established, we evaluated the impact of pharmacological IRE1 inhibition on the parameters mentioned above. However, a major hurdle in that process resided in the fact that MKC8866, an IRE1 inhibitor previously demonstrated to impact tumor growth in other cancer models [10] does not pass the blood brain barrier (BBB) and as such would not be suitable for intervention in brain tumors. To overcome this problem, we reasoned that we could take advantage of the surgical resection step for selective intra-operative delivery of IRE1-targeting drugs, provided that proper biomaterial scaffolds would be used to allow controlled release of the drugs to the tumor. The procedure was successfully set in place using two kinds of biomaterials including a drug containing fibrin microgel (conceived at NUI Galway) or a collagen sponge (Pangen) soaked in drug solutions (IRE1 RNase inhibitor MKC8866) at the appropriate concentrations (**Figure S1**). We followed up by evaluating how the implantation of these drug-containing devices impacted on tumor development (without Stupp-like treatment). Mouse survival was evaluated post implantation and revealed that local IRE1 inhibition did not significantly (neither positively nor negatively) modify mouse survival following resection (**Figure S1**). As we initially showed that IRE1 signaling and more particularly the IRE1/XBP1s axis contributed to the remodeling of the tumor stroma, we then sought to evaluate how the intra-operative application of the IRE1 RNase inhibitor impacted on tumor characteristics. To this end, we first evaluated tumor structures at sacrifice using H&E staining (**Figure S1B**). This showed that although the fibrin microgels appeared to exhibit a controlled tumor development (compared to non-resected tumors), implantation of the Pangen sponges did not modify (rather amplified) tumor development. The H&E analysis did not allow the characterization of other tumor properties. Collectively, implantation of the devices containing the IRE1 inhibitor did not modify mouse survival compared to surgery alone (**Figure S1C**). Following these observations, we decided to abandon the “sponge” approach as it was estimated very toxic. Moreover, the Pangen sponge was outperformed by the fibrin/collagen microgels regarding specific controlled release of the drug. As a consequence, we have since focused only on the fibrin/collagen gels and kept working with MKC8866 as this compound produced minimal toxicity (not shown).

To further document the impact of MKC8866-mediated inhibitory effects on GL261 cells, we performed *in vitro* experiments on these cells. We first evaluated whether MKC8866 blocked ER stress induced by tunicamycin (TUN, an inhibitor of protein N-glycosylation and inducer of ER stress); and then we evaluated the stress induced upon TMZ treatment. As expected MKC8866 treatment impaired TUN-induced IRE1 signals (**Figure 4A**), did not affect TUN-induced ATF4 expression (**Figure 4B**, top panel), but unexpectedly affected TMZ-induced ATF4 expression (**Figure 4B**, bottom panel). The activation of three UPR branches IRE1, PERK and ATF6 also appeared to be altered in response to TMZ (**Figure 4C, D**). Finally, since treatment of GL261 cells with MKC8866 appeared to impact on the response of these cells to different stressors, we evaluated the impact IRE1 RNase inhibition on cell sensitivity to TMZ. To do so, TMZ IC50 was determined for GL261 and evaluated to range to 800 μM (**Figure 4E**), and did not vary upon combined radiation. Remarkably, GL261 cells’ sensitivity to TMZ was increased upon treatment with MKC8866, phenomenon further amplified with combined irradiation (**Figure 4E**). These results suggest that TMZ-induced damages to the cells promote the activation of IRE1 dependent adaptive signals that if blocked by MKC8866, decrease the cell ability to cope with TMZ treatment. Consequently, these data suggest that combining Stupp treatment with MKC8866 might be an interesting approach to enhance treatment efficacy.

**Figure 4:**
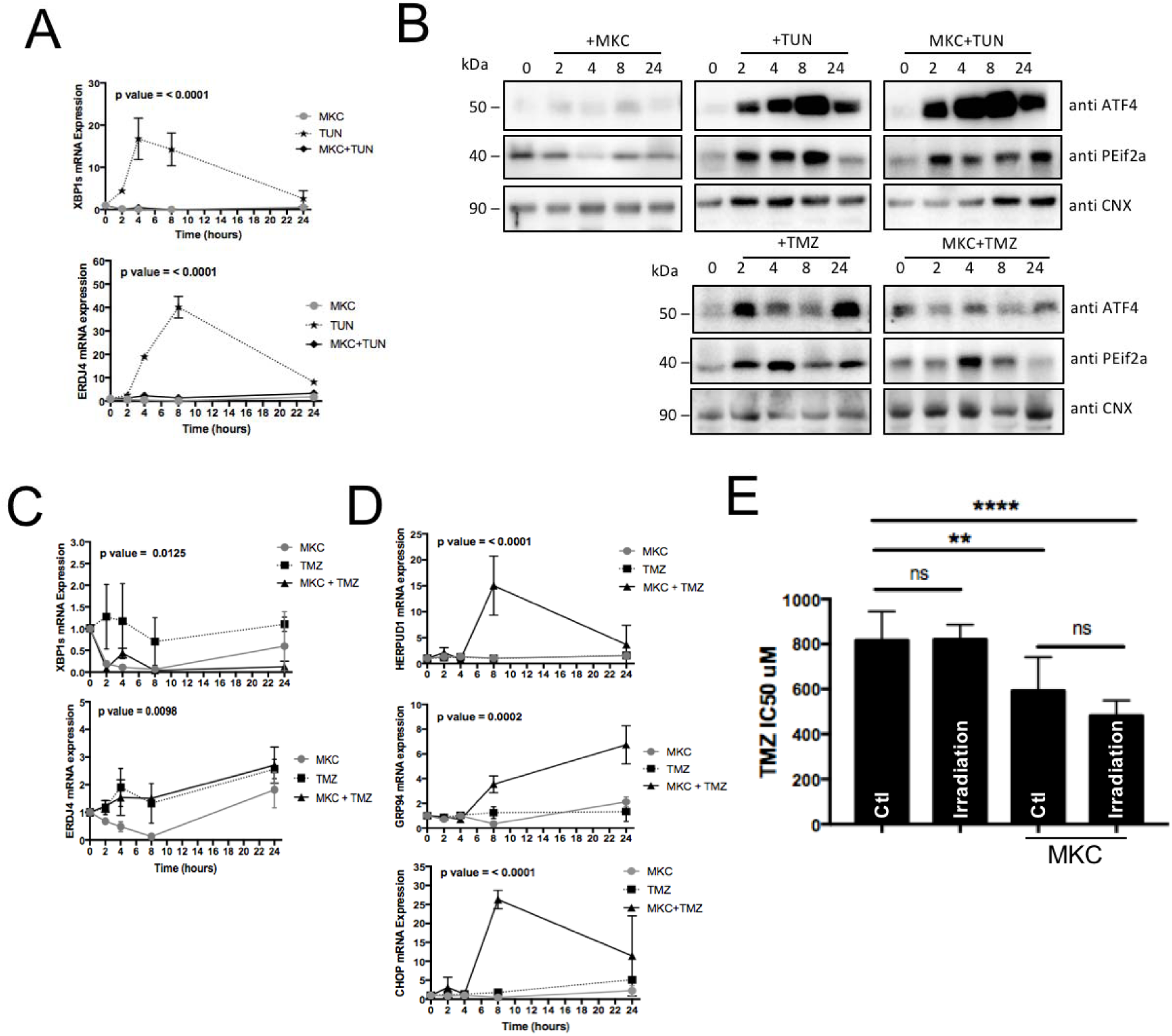
Evaluation of the impact of IRE1 inhibition by MKC8866 on GL261 response to stress. **A)** Impact of MKC8866 treatment on IRE1 signaling as monitored using RT-qPCR analysis of XBP1 mRNA splicing and expression of XBP1s target gene *Erdj4*, induced by tunicamycin. **B)** Impact of MKC8866 on TUN- and TMZ-induced Atf4 expression and the phosphorylation of eIF2α as monitored using Western blot. **C)** Impact of MKC8866 treatment on IRE1 signaling induced by TMZ as monitored using RT-qPCR analysis of XBP1 mRNA splicing and expression of XBP1s target gene *Erdj4*. **D)** Impact of MKC8866 treatment on ER stress signaling induced by TMZ as monitored using RT-qPCR analysis of HERPUD1, GRP94 and CHOP mRNA expression. **E)** Impact of MKC8866 treatment on the sensitivity of GL261 cells to TMZ treatment in the presence or absence of radiation.

To test this hypothesis, we evaluated the combination of MKC8866 with Stupp-like treatment using the fibrin-collagen gel plug as delivery tool. Surgery and drug implant were carried out and the first line of treatment including radiation/chemotherapy was done in the previous series of experiments (**Figure 5A**). Mouse survival was evaluated and brains were collected as sacrifice. In the first instance we evaluated the impact of MKC8866, Stupp-like treatment and combined therapy on tumor morphology at sacrifice. The new plugs of 2 mm^3^ were appropriate in terms of size and conformation, they adjusted perfectly to the resection cavity. Our latest results show that tumors developing in the presence of the MKC8866 containing plug and upon Stupp treatment present a totally different phenotype from those developing under Stupp treatment alone. This phenotype corresponds to much smaller tumors with tumor escape significantly delayed (**Figure 5B**). Furthermore, when necrosis was measured in tumors collected at sacrifice (H&E staining), we observed that the combination of Stupp treatment and MKC8866 yielded much important necrosis, reflecting treatment efficacy (**Figure 5C**). At last, we tested whether combined MKC8866 and Stupp treatment impacted on mouse survival. This allowed us to establish survival curves (**Figure 5D**) that demonstrate the positive impact of MKC8866 contained fibrin/collagen graft (1-2 μl) either at a 10 μM (**Figure 5D**, green) or 100 μM (**Figure 5D**, orange) concentration on the efficiency of Stupp treatment. We found that MKC8866 yielded a significant increase of mice survival (of about 20%) compared to Stupp alone however with no significant impact of the drug concentration used in the gel (**Figure 5D)** whereas it was observable in cell culture (**Figure S2**). This suggested that the main limitation in our experimental design was the capacity of the fibrin/collagen gel to control the delivery of the drug over time. Collectively, our results show that IRE1 RNase inhibition might represent an appealing approach to enhance the efficiency of current treatments towards GBM and that further work is needed to improve drug delivery to the tumor site.

**Figure 5:**
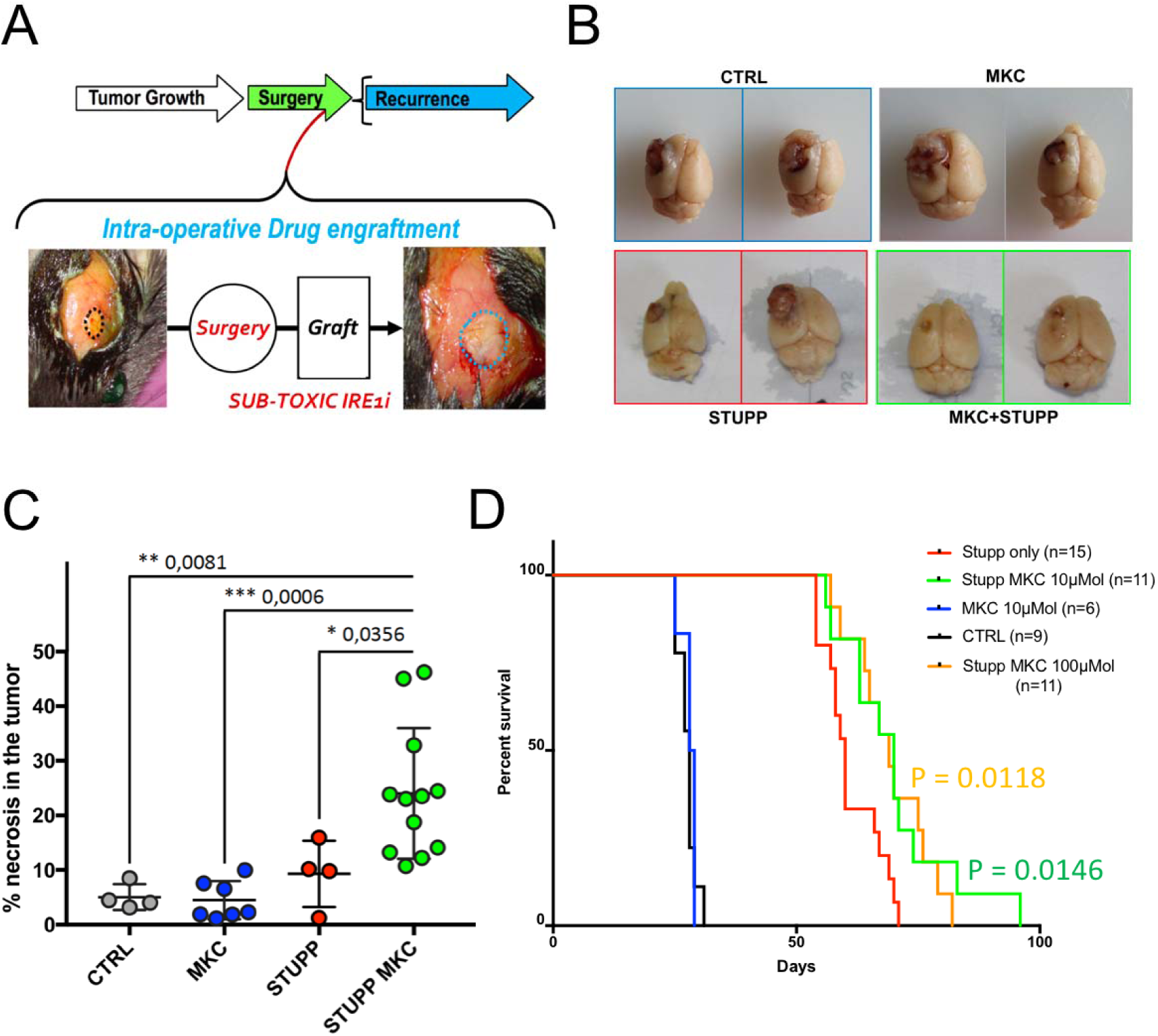
Impact of IRE1 inhibition on tumor development under Stupp treatment. **A)** Schematic representation of the experimental procedure and mouse survival. **B)** Macroscopic images of the brains following injection of GL261-Luc cells, surgical resection and treatment with ctrl plug (blue), MKC8866-containing fibrin microgels (grey); ctrl plug + Stupp (red) and MKC8866-containing fibrin microgels + Stupp (green). **C)** Quantification of tumor necrosis upon different treatments. **D)** Impact of IRE1 inhibition on tumor development under Stupp treatment - Kaplan-Meier representation. p value between Stupp and Stupp+MKC8866 (0.0146 - 10μM and 0.0118 - 100μM).

At last, three tumors from each group collected at sacrifice were dissociated and dissociated cells grown in culture to isolate GL261-derived lines following various treatments. These lines were then evaluated for their ability to proliferate (**Figure 6A**, bottom panel). This showed that except for cells derived from non-resected tumors which proliferated faster, none of the other lines showed any variation compared to the parental line, this result was similar to that observed with the sensitivity of cells to MKC8866 (**Figure 6A**, top panel). In contrast, all GL261-derived lines exhibited a higher sensitivity to TMZ than the parental line with the most striking observation with the lines derived from tumors relapsing after resection (**Figure 6A**, middle panel). It is interesting to note that lines derived from tumors treated with Stupp-like and MKC8866 showed great heterogeneity of response, a result contrasting with other lines. To investigate if this heterogeneity was reflected at the level of the transcriptome, lines were analyzed using RNAseq and then clustered based on IRE1 activity signature (**Figure 6B**). This revealed that treatment of the animals does not seem to impact on the basal IRE1 activity in tumor-derived GL261 cells grown in culture. This result might suggest that either the tumor cell selection process post-resection at sacrifice induces a strong bias in the nature of the cells collected, and in which case bulk tumor analyses would be necessary or that the alterations resulting from the treatment do not directly impact on basal IRE1 activity thereby suggesting that global functional analysis of these cells’ transcriptomes would be relevant to analyze further to identify the affected pathways.

**Figure 6:**
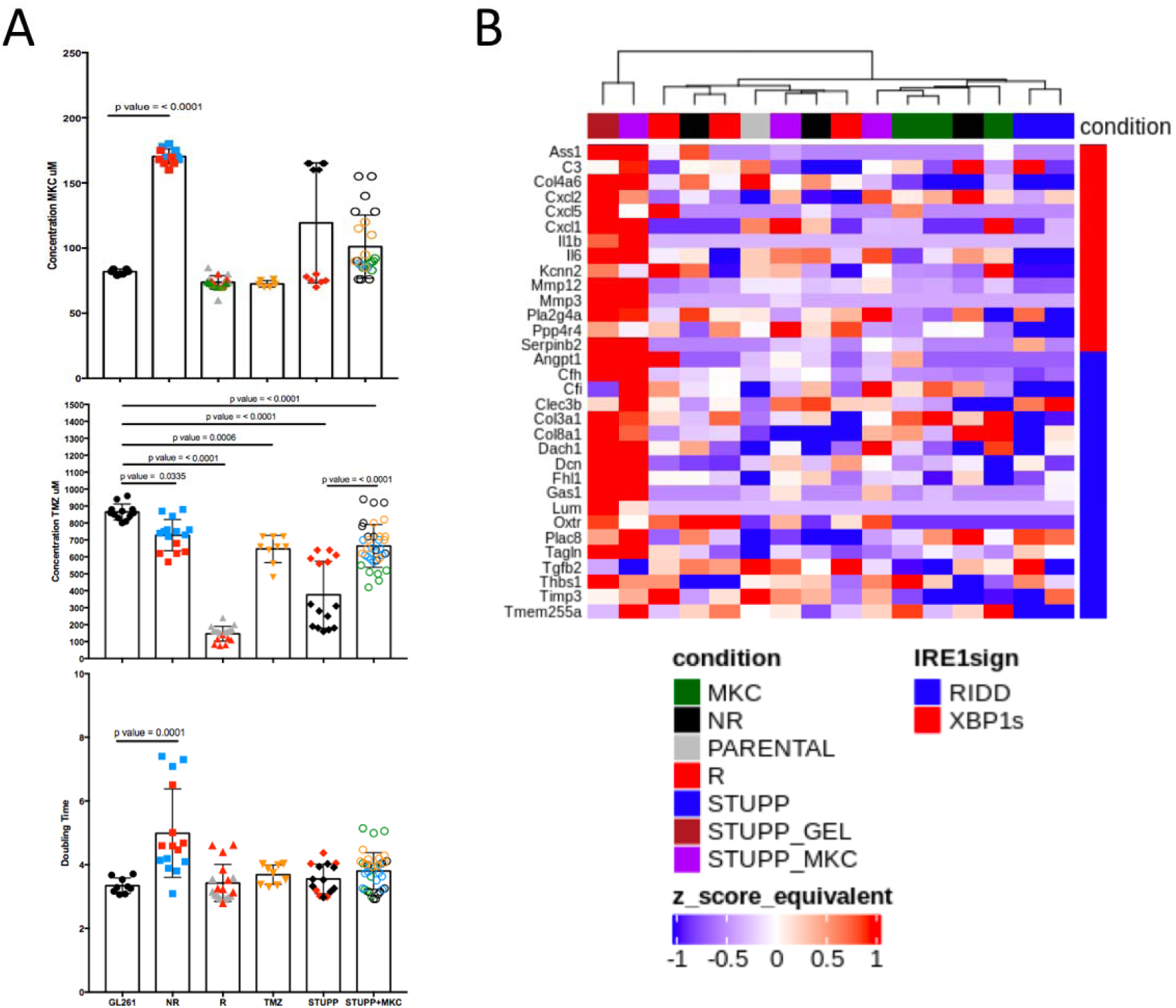
Analysis of GL261-derived lines following in vivo treatments. **A)** Characterization of the derived lines regarding their sensitivity to MKC8866 (top) or TMZ (middle), and regarding their proliferation ability. Twelve lines were analyzed in multiple experiments (n > 4). Shown is the average ±SD. **B)** RNAseq analysis of the different derived lines and representation as heat map. Hierarchical clustering was performed as described in the methods section.

## Discussion

Herein, we established a novel murine preclinical model of GBM that recapitulates standard therapy for human patients. This model includes surgical resection of the tumor and Stupp-like treatment combining irradiation and TMZ-based chemotherapy. Our model relies on the use of GL261 cells that exhibit a genome harboring the majority, if not all mutations, driving gliomagenesis. These mutations and/or copy number variations were observed in genes involved in the 3 main pathways affected in GBM (the receptor tyrosine kinase (RTK), P53, RB1 pathways, **Figure 1** [14]). Furthermore, GBM related genes, harboring one SNV in GL261 and two DNA copies (**Tables S1-S3**) had variable mutation allele frequency (MAF) values ranking between 6 and 100%. This MAF distribution strongly suggests that multiple, different clones compose the parental GL261 line (up to 5). Altogether these GL261 characteristics recapitulate molecular features observed for primary GBM at diagnostic with the presence of multiple drivers and/or multiple GBM clones [2]. Furtheremore we shown that GL261 displayed a constitutive activation of IRE1. Therefore, this cell line very likely constitutes a valuable GBM model for preclinical studies in mice, and in particular to study IRE1 inhibitors.

The use of such a model in the context of our discovery regarding the role of IRE1 signaling in GBM [15], has allowed us to test the impact of IRE1 pharmacological inhibition in enhancing the efficacy of Stupp treatment as already demonstrated in other contexts, for instance for the sensitization of Triple Negative Breast Cancers (TNBC) to paclitaxel therapy [10, 16]. In contrast with the approaches carried out with TNBC which included systemic injection of MKC8866 [10, 16], the available pharmacological inhibitors did not pass the blood brain barrier and as such we had to design a novel type of therapeutic strategy which would allow the controlled release of the drug during Stupp treatments. As such, we took advantage of the surgical step to use the peri-surgical implantation of a biocompatible graft containing the drug (**Figure S1**). This was achieved by using a fibrin/collagen graft (1-2 μl) containing 10 μM (**Figure 5D**, green) or 100 μM (**Figure 5D**, orange) of MKC8866, corresponding to non-toxic concentrations in cell culture and in animals. The grafts were implanted at the time of surgery and mice were then exposed to Stupp-like treatments. Both macroscopic exploration, immunohistochemistry and Kaplan-Meier survival curves (**Figure 5**) indicated that combination of MKC8866 with Stupp treatment enhanced its efficacy, leading to increased necrosis within the tumor and to prolonged mouse survival.

Collectively, our work provides the proof of concept that the pharmacological inhibition of IRE1 could be used as an adjuvant to enhance the efficacy of combined radiation and chemotherapy in the context of GBM. However, one still must evaluate the impact of these treatments on the characteristics of the recurrent tumors regarding their treatment resistance properties. Also of note is to investigate whether specific strata from cohorts of GBM patients (e.g., depending on their IRE1 signaling characteristics) would derive more benefit and should be preferentially targeted.

## Acknowledgements

We thank the BIOSIT H2P2 platform and Florence Jouan for immunohistochemistry and the BIOSIT Animal facility ARCHE (https://biosit.univ-rennes1.fr/). This work was funded by grants from the French National Cancer Institute (INCa, PLBIO), the Fondation pour la recherche Médicale (FRM; équipe labellisée 2018) to EC and EU H2020 MSCA ITN-675448 (TRAINERS) and MSCA RISE-734749 (INSPIRED) grants to AS, AC and EC. The work was also supported in part by research grant from Science Foundation Ireland (SFI), co-funded under the European Regional Development Fund under Grant number 13/RC/2073.

## Authors’ contributions

**Conceptualization** – PJLR, RP, EC; **Data curation** – MA, NS, KV, AE, JM; **Formal analysis** – MA, NS; **Funding acquisition** – EC, AP, AS, AG; **Investigation** – PJLR, KV, RP, GJ, SL, MA, NS, AE, JM, TA; **Methodology** – PJLR, RP, KV, EC; **Project administration** – PJLR, RP, TA, EC; **Resources** – JS, AP, JBP, AC, QZ; **Supervision** – EC; **Validation** – PJLR, RP, EC; **Visualization** – PJLR, RP, KV, GJ, SL, MA, NS, AE, AC, JM, EC; **Writing – original draft** – EC; **Writing – review & editing** – PJLR, RP, SL, AG, AS, MA, NS, AE, JM, AC, TA, EC (established according to https://casrai.org/credit/)

