## Supplementary material for "Local intracerebral Inhibition of IRE1 by MKC8866 sensitizes glioblastoma to irradiation/chemotherapy *in vivo*": Suppl material

^1^Inserm U1242, University of Rennes, Rennes, France. ^2^Centre de lutte contre le cancer Eugène Marquis, Rennes, France. ^3^Rennes Brain Cancer Team (REACT), 35000 Rennes, France.^4^Neurosurgery dept, University Hospital of Rennes, 35000 Rennes, France.^5^Institute of Chemical Biology, National Hellenic Research Foundation (N.H.R.F.), Athens, Greece. ^6^Department of Biochemistry and Biotechnology, University of Thessaly, Viopolis, 41500, Larissa, Greece; ^7^CÚRAM, Centre for Research in Medical Devices, National University of Ireland, Galway, Ireland.^8^Apoptosis Research Centre, National University Ireland Galway, Galway, Ireland.^9^Fosun OrinovePharmaTech Inc., 3537 Old Conejo Road, Suite 104, Newbury Park, CA, 91320, USA.^10^University of Rennes, CNRS, IGDR [(Institut de génétique et développement de Rennes)]-UMR 6290, F-35000 Rennes, France.^11^CHU Rennes, Service de Génétique Moléculaire et Génomique Médicale, Rennes, France.^12^e-NIOS PC, Kallithea-Athens, Greece.^13^University of Rennes, Plateforme GEH, CNRS, Inserm, BIOSIT - UMS 3480, US_S 018, F-35000 Rennes, France.

**SUPPLEMENTAL MATERIAL**

Table S1

Table S2

Table S3

Movie S1

Figure S1

Figure S2

**Table S1**: **GBM related genes** from Lee et al. Nature 2018 (blue) and additional genes (red). Mouse homologs were retrieved from Ensembl.

| ***hgnc_symbol*** | ***mmusculus_homolog_associated_gene_name*** | **chromosome** | **start** | **end** | **strand** |
| --- | --- | --- | --- | --- | --- |
| *ABCB1* | *Abcb1a* | 5 | 8660077 | 8748575 | + |
| *ABCB1* | *Abcb1b* | 5 | 8798147 | 8866315 | + |
| *ABCC9* | *Abcc9* | 6 | 142587862 | 142702315 | - |
| *ADAM29* | *Adam29* | 8 | 55870912 | 55906948 | - |
| *AFM* | *Afm* | 5 | 90518932 | 90553543 | + |
| *ANKRD36* | *Ankrd36* | 11 | 5569697 | 5660839 | + |
| *ARID1A* | *Arid1a* | 4 | 133679008 | 133756769 | - |
| *ARID1B* | *Arid1b* | 17 | 4994332 | 5347656 | + |
| *ATF6* | *Atf6* | 1 | 170704674 | 170867771 | - |
| *ATRX* | *Atrx* | X | 105797615 | 105929403 | - |
| *BRAF* | *Braf* | 6 | 39603237 | 39725463 | - |
| *C1orf150/GCSAML* |  |  |  |  |  |
| *CALCR* | *Calcr* | 6 | 3685680 | 3764714 | - |
| *CARD6* | *Card6* | 15 | 5095981 | 5108539 | - |
| *CD3EAP* | *Cd3eap* | 7 | 19356014 | 19359483 | - |
| *CDH18* | *Cdh18* | 15 | 22549022 | 23474418 | + |
| *CDH9* | *Cdh9* | 15 | 16728756 | 16857094 | + |
| *CDHR3* | *Cdhr3* | 12 | 33033796 | 33092875 | - |
| *CDK4* | *Cdk4* | 10 | 127063534 | 127067920 | + |
| *CDK6* | *Cdk6* | 5 | 3341485 | 3531008 | + |
| *CDKN2A* | *Cdkn2a* | 4 | 89274471 | 89294653 | - |
| *CDKN2B* | *Cdkn2b* | 4 | 89306299 | 89311039 | - |
| *CDKN2C* | *Cdkn2c* | 4 | 109660876 | 109667189 | - |
| *CDX4* | *Cdx4* | X | 103321398 | 103330592 | + |
| *CIC* | *Cic* | 7 | 25267704 | 25294159 | + |
| *COL1A2* | *Col1a2* | 6 | 4504814 | 4541544 | + |
| *CXorf22/CFAP47* |  |  |  |  |  |
| *DCAF12L2* | *Dcaf12l2* | X | 44365458 | 44368342 | - |
| *DCAF12L2* | *Dcaf12l1* | X | 44786570 | 44790197 | - |
| *DGKI* | *Dgki* | 6 | 36846022 | 37300184 | - |
| *DRD5* | *Drd5* | 5 | 38319367 | 38322518 | + |
| *DYNC1I1* | *Dync1i1* | 6 | 5725639 | 6028039 | + |
| *EGFR* | *Egfr* | 11 | 16752203 | 16918158 | + |
| *EIF2AK3* | *Eif2ak3* | 6 | 70844515 | 70905245 | + |
| *ERN1* | *Ern1* | 11 | 106397620 | 106487796 | - |
| *FGA* | *Fga* | 3 | 83026076 | 83033627 | + |
| *FGFR/FGFR1OP2* | *Fgf17-201* | 14 | 70636203 | 70642268 | - |
| *FOXR2* | *Foxr2* | X | 153118786 | 153132861 | + |
| *FRMD7* | *Frmd7* | X | 50895180 | 50942710 | - |
| *FUBP1* | *Fubp1* | 3 | 152210422 | 152236826 | + |
| *GABRA1* | *Gabra1* | 11 | 42130939 | 42182930 | - |
| *GABRA6* | *Gabra6* | 11 | 42306437 | 42321072 | - |
| *GABRB2* | *Gabrb2* | 11 | 42419757 | 42629028 | + |
| *GPX5* | *Gpx5* | 13 | 21286429 | 21292731 | - |
| *IDH1* | *Idh1* | 1 | 65158616 | 65186500 | - |
| *IDH2* | *Idh2* | 7 | 80094846 | 80115392 | - |
| *IL18RAP* | *Il18rap* | 1 | 40515362 | 40551705 | + |
| *KEL* | *Kel* | 6 | 41686330 | 41704339 | - |
| *KEL* | *Kel* | 6 | 41686330 | 41704339 | - |
| *KRAS* | *Kras* | 6 | 145216699 | 145250239 | - |
| *KRTAP20-2* | *Krtap20-2* | 16 | 89205861 | 89206388 | - |
| *LCE4A* |  |  |  |  |  |
| *LRRC55* | *Lrrc55* | 2 | 85162334 | 85196699 | - |
| *LUM* | *Lum* | 10 | 97565128 | 97572703 | + |
| *LZTR1* | *Lztr1* | 16 | 17508688 | 17526333 | + |
| *MDM2* | *Mdm2* | 10 | 117688875 | 117710758 | - |
| *MDM4* | *Mdm4* | 1 | 132959484 | 133030561 | - |
| *MET* | *Met* | 6 | 17463800 | 17573980 | + |
| *MGMT* | *Mgmt* | 7 | 136894614 | 137130266 | + |
| *MLL2/KMT2B* | *Kmt2b* | 7 | 30568858 | 30588726 | - |
| *MMP13* | *Mmp13* | 9 | 7272514 | 7283331 | + |
| *NF1* | *Nf1* | 11 | 79339693 | 79581612 | + |
| *NLRP5* | *Nlrp5* | 7 | 23385889 | 23441922 | + |
| *NOTCH1* | *Notch1* | 2 | 26457903 | 26516663 | - |
| *NOTCH2* | *Notch2* | 3 | 98013527 | 98150361 | + |
| *NRAS* | *Nras* | 3 | 103058285 | 103067914 | + |
| *ODF4* | *Odf4* | 11 | 68921835 | 68927081 | - |
| *PARD6B* | *Pard6b* | 2 | 168081004 | 168101203 | + |
| *PDGFRA* | *Pdgfra* | 5 | 75152292 | 75198215 | + |
| *PIK3CA* | *Pik3ca* | 3 | 32397671 | 32468486 | + |
| *PIK3R1* | *Pik3r1* | 13 | 101680563 | 101768217 | - |
| *PLCH2* | *Plch2* | 4 | 154983115 | 155056784 | - |
| *PLCH2* | *Plch2* | 4 | 154983115 | 155056784 | - |
| *PODNL1* | *Podnl1* | 8 | 84125989 | 84132527 | + |
| *PTEN* | *Pten* | 19 | 32757497 | 32826160 | + |
| *PTEN* | *Pten* | 19 | 32757497 | 32826160 | + |
| *QKI* | *Qk* | 17 | 10202601 | 10319854 | - |
| *RB1* | *Rb1* | 14 | 73183673 | 73325822 | - |
| *RFX6* | *Rfx6* | 10 | 51677756 | 51730432 | + |
| *RPL5* | *Rpl5* | 5 | 107900502 | 107909005 | + |
| *SCN9A* | *Scn9a* | 2 | 66480080 | 66634962 | - |
| *SEMA3C* | *Sema3c* | 5 | 17574281 | 17730268 | + |
| *SEMA3E* | *Sema3e* | 5 | 14025276 | 14256689 | + |
| *SEMG1* | *Svs2* | 2 | 164235929 | 164238466 | - |
| *SEMG1* | *Svs3a* | 2 | 164289268 | 164291500 | + |
| *SEMG1* | *Svs3b* | 2 | 164254363 | 164256640 | - |
| *SETD2* | *Setd2* | 9 | 110532597 | 110618633 | + |
| *SIGLEC8* | *Siglece* | 7 | 43651070 | 43660161 | - |
| *SMARCA4* | *Smarca4* | 9 | 21616169 | 21704230 | + |
| *TERT* | *Tert* | 13 | 73626911 | 73649843 | + |
| *TP53* | *Trp53* | 11 | 69580359 | 69591873 | + |

**Table S2: GL261 cell line observed variants for GBM related genes.** The predicted functional impact on the transcript is given by the Ensembl Variant Effect Predictor algorithm. A subjective classification of the severity of the variant consequence (HIGH, MODERATE or LOW) is given for compatibility with other variant annotation tools.

| **IMPACT** | **Effect on transcript** | **Nb of variants** |
| --- | --- | --- |
| HIGH | stop_gained | 2 |
| HIGH | frameshift_variant | 1 |
| MODERATE | missense_variant | 19 |
| LOW | synonymous_variant | 4 |
| LOW | splice_region_variant&intron_variant | 2 |
| UNKNOWN | intron_variant | 1617 |
| UNKNOWN | downstream_gene_variant | 39 |
| UNKNOWN | 3_prime_UTR_variant | 28 |
| UNKNOWN | upstream_gene_variant | 26 |
| UNKNOWN | 5_prime_UTR_variant | 12 |
| UNKNOWN | intron_variant&non_coding_transcript_variant | 6 |
| UNKNOWN | intergenic_variant | 6 |
| UNKNOWN | regulatory_region_variant | 1 |

**Table S3: Copy-Number Variants and allelic imbalances for GBM related genes in GL261.**

| **Symbol** | **Chromosome** | **CNV_Start** | **CNV_Stop** | **Copy nb** | **ALT** |
| --- | --- | --- | --- | --- | --- |
| Adam29 | 8 | 52333775 | 71668335 | 3 | gain |
| Afm | 5 | 67477105 | 92807020 | 3 | gain |
| Ankrd36 | 11 | 0 | 24053585 | 3 | gain |
| Arid1A | 4 | 132841615 | 133887300 | 1 | loss |
| Calcr | 6 | 0 | 14217920 | 1 | loss |
| Card6 | 15 | 0 | 10708720 | 6 | gain |
| Cdh9 | 15 | 10708720 | 21033975 | 7 | gain |
| Cdh18 | 15 | 21672140 | 27885405 | 6 | gain |
| Cdk4 | 10 | 126608540 | 129113090 | 3 | gain |
| Cdkn2c | 4 | 107422555 | 117304915 | 1 | loss |
| Col1a2 | 6 | 0 | 14217920 | 1 | loss |
| Dcaf12l1 | X | 34609485 | 74412020 | 1 | loss |
| Dcaf12l2 | X | 34609485 | 74412020 | 1 | loss |
| Drd5 | 5 | 32740270 | 50978205 | 3 | gain |
| Dync1i1 | 6 | 0 | 14217920 | 1 | loss |
| Egfr | 11 | 0 | 24053585 | 3 | gain |
| Emx2 | 19 | 51053200 | 60629920 | 1 | loss |
| Foxr2 | X | 152201645 | 157855985 | 3 | gain |
| Frmd7 | X | 34609485 | 74412020 | 1 | loss |
| Gabra1 | 11 | 37447975 | 63154280 | 3 | gain |
| Gabra6 | 11 | 37447975 | 63154280 | 3 | gain |
| Gabrb2 | 11 | 37447975 | 63154280 | 3 | gain |
| ERN1 (IRE1) | 11 | 101120145 | 108292780 | 4 | gain |
| Lum | 10 | 74005915 | 117711020 | 3 | gain |
| Mdm2 | 10 | 74005915 | 117711020 | 3 | gain |
| Nf1 | 11 | 74799730 | 80488030 | 3 | gain |
| Pdgfra | 5 | 67477105 | 92807020 | 3 | gain |
| Pik3r1 | 13 | 90101540 | 116320075 | 1 | loss |
| Plch2 | 4 | 138904890 | 156508905 | 1 | loss |
| Pten | 19 | 21839110 | 42478300 | 1 | loss |
| Rfx6 | 10 | 32869035 | 68944460 | 3 | gain |
| Sema3c | 5 | 15831020 | 19679820 | 3 | gain |
| Setd2 | 9 | 110174730 | 111278430 | 1 | loss |
| Tert | 13 | 66488020 | 86357450 | 1 | loss |

**Movie S1: GBM surgical resection procedure.**

**Figure S1:** **Development of an intra-operative strategy for IRE1 inhibitors delivery to the site of tumor resection.** This includes the development of specific biomaterial scaffolds allowing controlled release of the drugs. **A)** Schematic representation of the experimental design. **B)** Impact of IRE1 inhibition *in vivo*. Schematic representation of the experiment (top) together with H&E staining of the resulting tumors at sacrifice (bottom). **C)** Kaplan-Meier representation of mouse survival under this regiment (black: non-resected, red: surgical resection, blue/green: IRE1 inhibitor, orange: surgical resection + radio/chemotherapy).

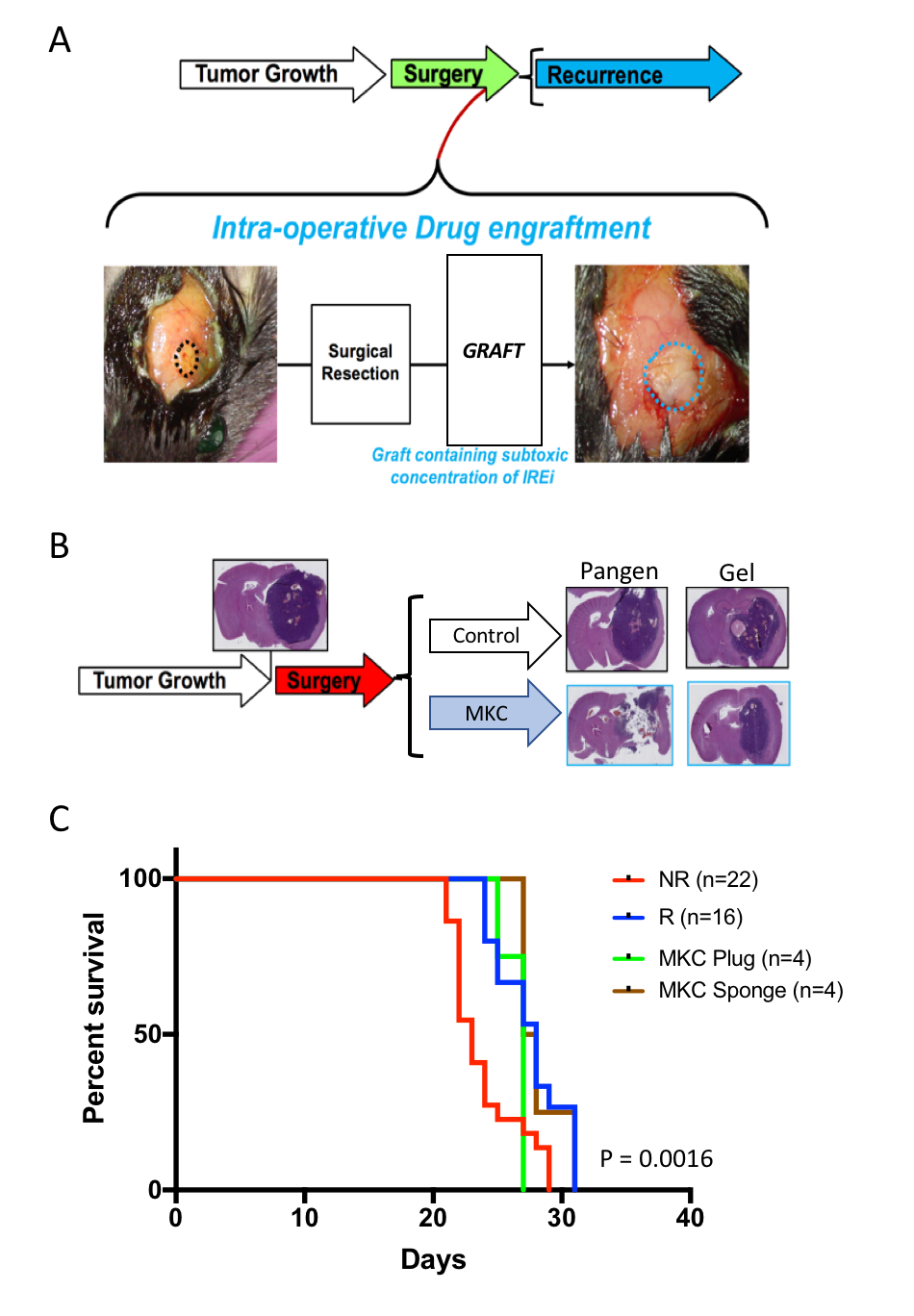

**Figure S2:** **MKC8866-containing fibrin microgels inhibit IRE1-mediated XBP1 splicing in the human glioblastoma cell lines U251.** U251 cells were cultivated in absence (no gel) or presence of fibrin microgels containing different concentrations of MKC8866 (0 to 100 µM, left and right panels) and different concentration of fibrin (30 or 60 mg/mL, right panel). Measurement of XBP1 splicing with or without tunicamycin induction of IRE1 by RT-PCR shows a significant inhibition of XBP1 splicing when the cells are cultivated with the microgels containing MKC8866, indicating a stable release of MKC8866 over one week. Increased concentration of MKC8866 potentiates this XBP1 splicing inhibition, whereas the fibrin concentration seems not to affect significantly the XBP1 splicing.

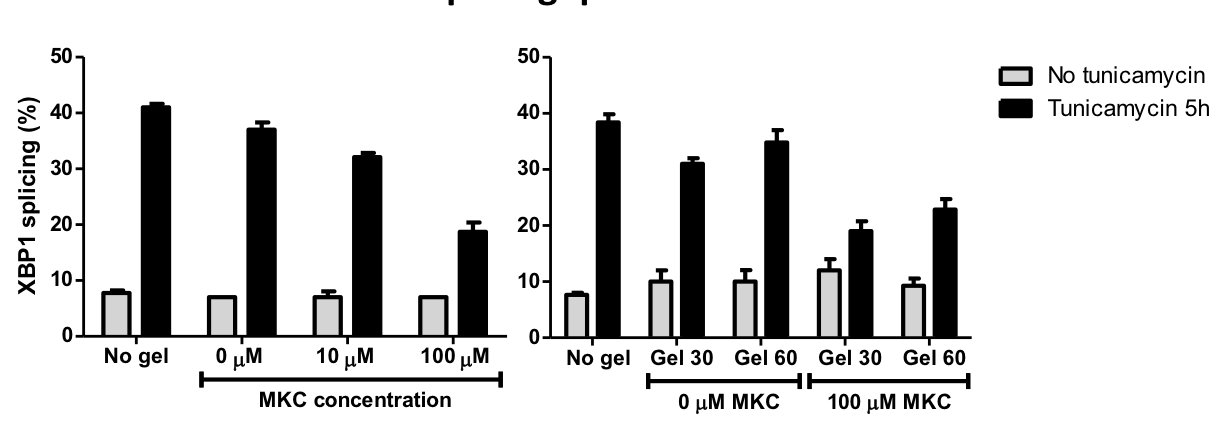
